# Foxm1 regulates neuronal progenitor fate during spinal cord regeneration

**DOI:** 10.1101/2020.02.26.962977

**Authors:** Diane Pelzer, Lauren S. Phipps, Raphael Thuret, Syed Murtuza Baker, Karel Dorey

## Abstract

Mammals have limited tissue regeneration capabilities, particularly in the case of the central nervous system. Spinal cord injuries are often irreversible and lead to the loss of motor and sensory function below the site of the damage [1]. In contrast, amphibians such as *Xenopus* tadpoles can regenerate a fully functional tail, including their spinal cord, following amputation [2,3]. A hallmark of spinal cord regeneration is the re-activation of Sox2/3+ progenitor cells to promote regrowth of the spinal cord and the generation of new neurons [4,5]. In axolotls, this increase in proliferation is tightly regulated as progenitors switch from a neurogenic to a proliferative division via the planar polarity pathway (PCP) [6–8]. How the balance between self-renewal and differentiation is controlled during regeneration is not well understood. Here, we took an unbiased approach to identify regulators of the cell cycle expressed specifically in *X.tropicalis* spinal cord after tail amputation by RNAseq. This led to the identification of Foxm1 as a potential key transcription factor for spinal cord regeneration. *Foxm1*-/- *X.tropicalis* tadpoles develop normally but cannot regenerate their spinal cords. Using single cell RNAseq and immunolabelling, we show that *foxm1*+ cells in the regenerating spinal cord undergo a transient but dramatic change in the relative length of the different phases of the cell cycle, suggesting a change in their ability to differentiate. Indeed, we show that Foxm1 does not regulate the rate of progenitor proliferation but is required for neuronal differentiation leading to successful spinal cord regeneration.

## Results

### Foxm1 is specifically expressed in the regenerating spinal cord

We compared the transcriptome of isolated spinal cords at 1day post amputation (1dpa) and 3dpa to spinal cord from intact tails (0dpa, Figure 1A). Principle component plot, dendogram of sample-to-sample distances and MA-plot of the log fold change (FC) of expression in relation to the average count confirmed the quality of the data (Figures S1A-D). Between 0dpa and 1dpa, 5129 differentially expressed (DE) transcripts (FC> 2 and FDR<0.01) were identified (2074 down-, 3055 up-regulated). Between 0dpa and 3dpa, 9787 genes are differentially expressed (4609 down and 5178 up-regulated, Figure S1E).

**Figure 1.**
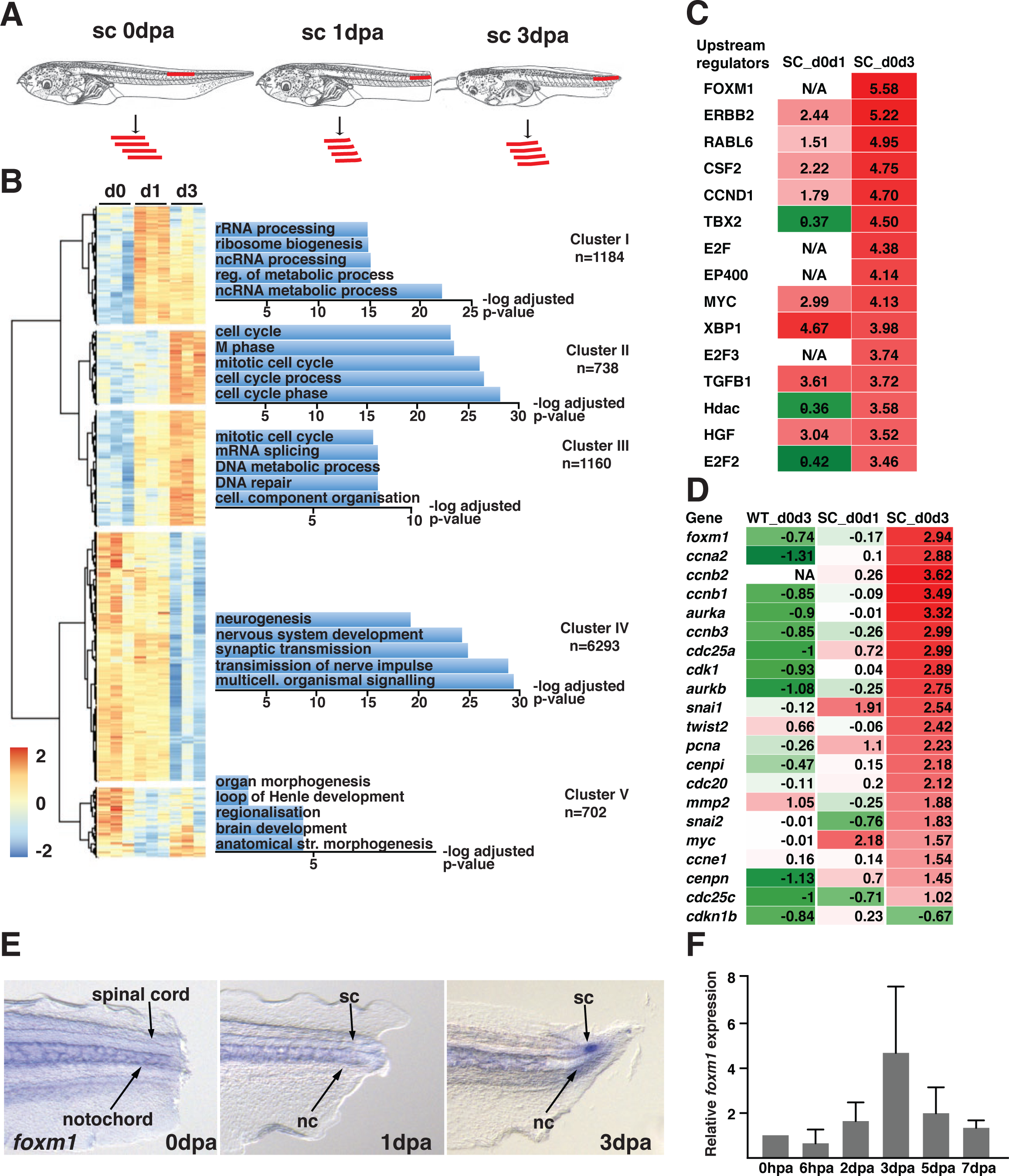
Analysis of differentially expressed genes during spinal cord regeneration. **(A)** Twenty spinal cords of NF50 tadpoles were isolated at 0, 1 and 3 day post amputation (dpa) and pooled for RNA sequencing **(B)** Genes with a |log2(FC)|>1 and p-adj<0.01 were used for hierarchical clustering. For each cluster, the gene list was uploaded on Fidea (http://circe.med.uniroma1.it/fidea/) [31]. The five most significant enrichment of GO (biological processes) terms are shown, and the -log10 (with Bonferroni correction) is shown. **(C)** The dataset was uploaded on the Ingenuity Pathway Analysis software (Qiagen). Genes with a |log2(FC)|>1 and p_adj<0.001 were considered. The software then identified upstream regulators based on the changes in expression levels of known dowstream target. Each upstream regulator is attributed a z-score, corresponding to the negative log of the p-value derived from the Fisher’s Exact test. **(D)** Changes in the expression of *foxm1* and known downstream targets in the whole tail comparing day 0 and day 3 (WT_d0d3) and in the spinal cord comparing day 0 and day 1(SC_d0d1) and day 0 and day 3 (SC_d0d3). The whole tail dataset was obtained from [9]. **(E)** Tadpoles at NF50 were amputated, fixed at the indicated time and then processed for WISH using a probe specific for *foxm1*. **(F)** Total RNA was isolated from regenerating tails at indicated time points post amputation, reversed transcribed into cDNA and analysed for *foxm1* expression by qPCR, using *ef1*α as a reference gene (n=3). The graph represents**(A)** Tadpoles at NF50 were amputated, fixed at the indicated time and then processed for WISH using a probe specific for *foxm1*. **(B)** Total RNA was isolated from regenerating tails at indicated time points post amputation, reversed transcribed into cDNA and analysed for *foxm1* expression by qPCR, using *ef1*α as a reference gene (n=3) the mean +/- SD of three independent experiments.

To identify the most enriched biological processes by gene ontology (GO), a non-biased hierarchical cluster for all DE genes was performed (Figure 1B). We observed three phases: first an increase in expression of genes involved in metabolic processes (cluster I), then a strong upregulation of genes associated with cell cycle regulation (cluster II and III) and finally, a downregulation of expression of genes involved in nervous system development (Cluster IV and V, Figure 1B).

Using Ingenuity Pathway Analysis (IPA), we identified potential upstream regulators that could explain changes in expression of downstream target genes, with Foxm1 showing the highest significance at 3dpa (Figure 1C). Using published RNAseq of tail regeneration in *X.tropicalis* [9], we compared changes in expression of known Foxm1 target genes between 0 and 3dpa in whole tail (WT_d0d3), 0 and 1dpa (SC_d0d1) and 0 and 3dpa (SC_d0d3) in spinal cord. *Foxm1* and its transcriptional targets are significantly upregulated only in the spinal cord at 3dpa, but not in the whole tail (Figure 1D).

We wanted to confirm the expression of *foxm1* during regeneration by *in situ* hybridisation (ISH) and RT-qPCR. ISH shows that *foxm1* is not expressed in the spinal cord at 0 and 1dpa but is restricted to the regenerating spinal cord at 3dpa (Figure 1E). We then performed RT-qPCR for *foxm1* over a period of 7 days, its expression peaks at 3dpa and decreases back to baseline levels at 7dpa (Figure 1F).

We next wanted to identify the upstream signal(s) regulating its expression. As *foxm1* expression starts at 3dpa, it is not a direct response to the injury. We tested if signalling pathways required for tail regeneration promote *foxm1* expression at 3dpa. A sustained increase of reactive oxygen species (ROS) in the tail is required for its regeneration [10]. ROS levels were decreased following amputation using DPI, an inhibitor of the NADPH Oxidase (NOX). In NF50 tadpoles treated with DPI from 36hpa until 72hpa, *foxm1* expression decreases by 69% (p=0.032) compared to DMSO controls (Figures S1F and S1G). ROS are upstream of different signalling pathways, including FGF [10,11]. Furthermore, Sonic hedgehog (Shh) signalling is also required for tail regeneration [12] and induces *Foxm1* expression in the developing cerebellar granule neuron precursors [13]. Amputated tails treated with an FGF receptor kinase inhibitor (SU5402, Figure S1H), or a Shh signalling inhibitor (cyclopamine, Figure S1I) failed to regenerate (data not shown) but no changes in *foxm1* expression was observed.

### Foxm1 is required for spinal cord regeneration

To test the role of Foxm1 during regeneration, we designed a guide RNA (gRNA) targeted at bases 129-152 downstream of the ATG to knockdown and knockout *foxm1* expression using CRISPR/Cas9 (Figure S2A). The efficacy of the gRNA was assessed by Restriction Fragment Length Polymorphism analysis. Co-injection of the gRNA with *cas9* mRNA did not lead to the destruction of the NcoI site but co-injection with 0.6ng and 1.5ng of Cas9 protein leads to NcoI-resistant PCR products in dose-dependent fashion (Figure S2B). We then tested the level of indels by sequencing individual clones leading to 50 to 90% mutation on the *foxm1* locus (Figure S2C). Injection of CRISPR/Cas9 injection at 1-cell stage leads to a 52% reduction of *foxm1* expression (p=0.0251; Figure S2D).

We then compared the ability of NF50 tadpoles injected with Cas9 alone (control) and Cas9 + *foxm1* gRNA (*foxm1* KD) to regenerate their spinal cord and tail. The ratio of regeneration was determined between 3 and 9dpa by dividing the length of the regenerating spinal cord by the length of the amputated spinal cord. No differences were observed at 3dpa, but from 5 to 9dpa the rate of regeneration was on average 40% lower (p=0.04) compared to controls (Figures 2A-C).

**Figure 2.**
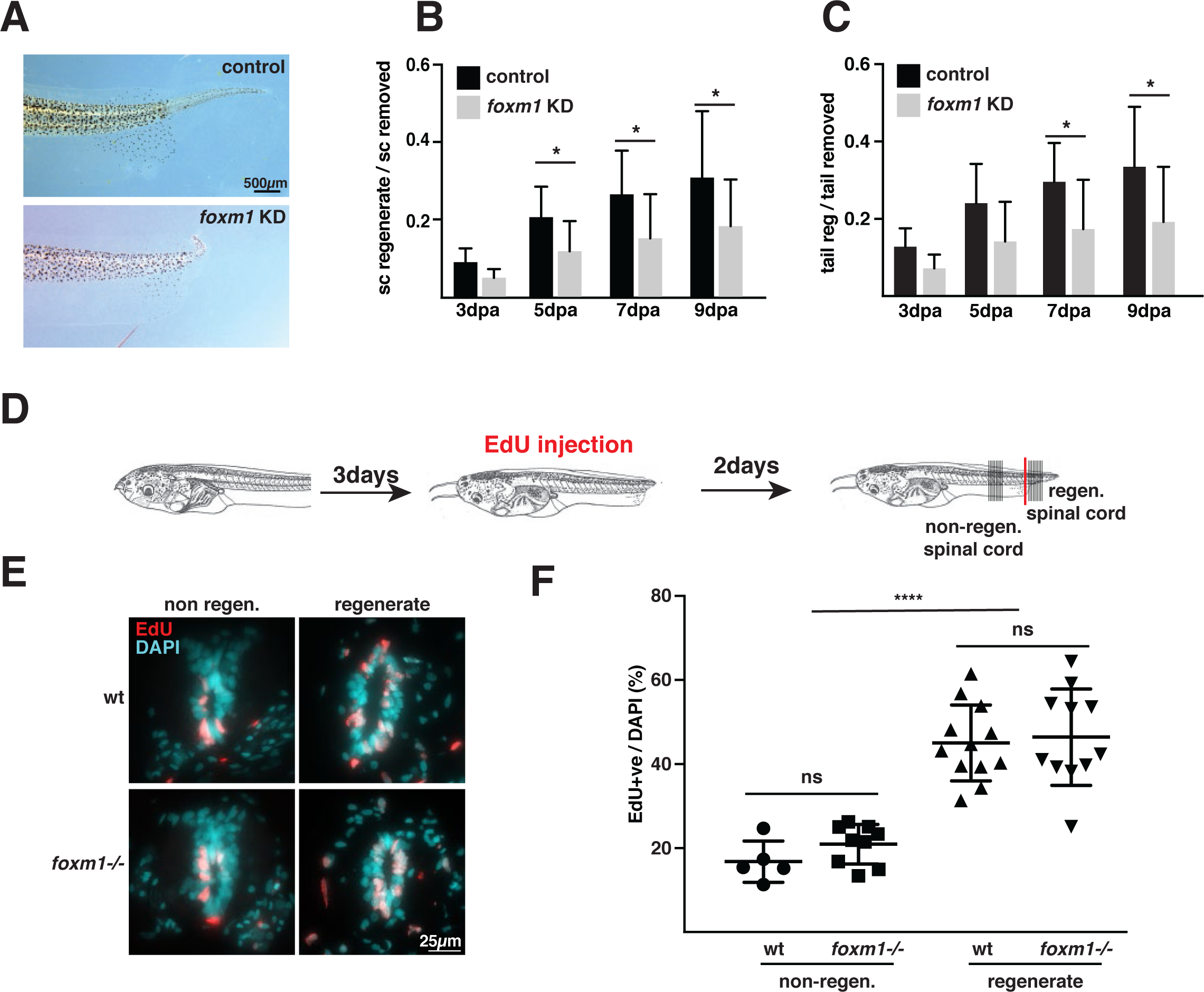
Foxm1 is required for spinal cord regeneration but does not regulate the length of the cell cycle. **(A)** A third of the tails of *foxm1* knockdown and wt tadpoles at NF50 were amputated and the tadpoles left to regenerate for 9 days. The images show representative tails at 9dpa. **(B-C)** To quantify the rate of regeneration, the ratio of the length of the regenerate to the length that has originally be amputated was compared for the spinal cord (B) and the whole tail (C). The graph represents the mean+/-SD of 3 independent experiments with at least 5 tadpoles in each experiment. **(D)** Experimental set up for EdU labelling, *foxm1* knockout and wt tadpoles were amputated and left to regenerate for 3 days. Tadpoles were then injected with EdU and 2 days later the tails were fixed and stained for EdU and DAPI. **(E)** Representative images of EdU (red) and DAPI (blue) staining **(F)** The graph represents the proportion of EdU+ cells over the total number of cells in the spinal cord. Each data point represents an embryo (mean of 8 to 10 sections +/- SD) and 5 to 12 embryos were analysed. *p<0.05 **** p<0.001, ns: non specific.

Could the impaired regeneration be caused by defective proliferation? To determine the rate of proliferation in the regenerating spinal cord, wt and *foxm1-/-* tadpoles were injected with EdU at 3dpa, followed by a 2-day chase (Figures 2D-F). As expected, we observed a higher proportion of EdU+ cells in the regenerate than in the non-regenerating spinal cord (∼45% and ∼20% respectively). However, no difference in EdU+ cells between wt and *foxm1*-/- was observed, showing that Foxm1 does not affect the overall length of cell cycle.

### Characterisation of *foxm1*+ cells during regeneration

To understand the role of Foxm1 during spinal cord regeneration, we characterised this cell population at the molecular level using single cell RNA sequencing (scRNA-seq) (Figures 3A and S3A).

**Figure 3.**
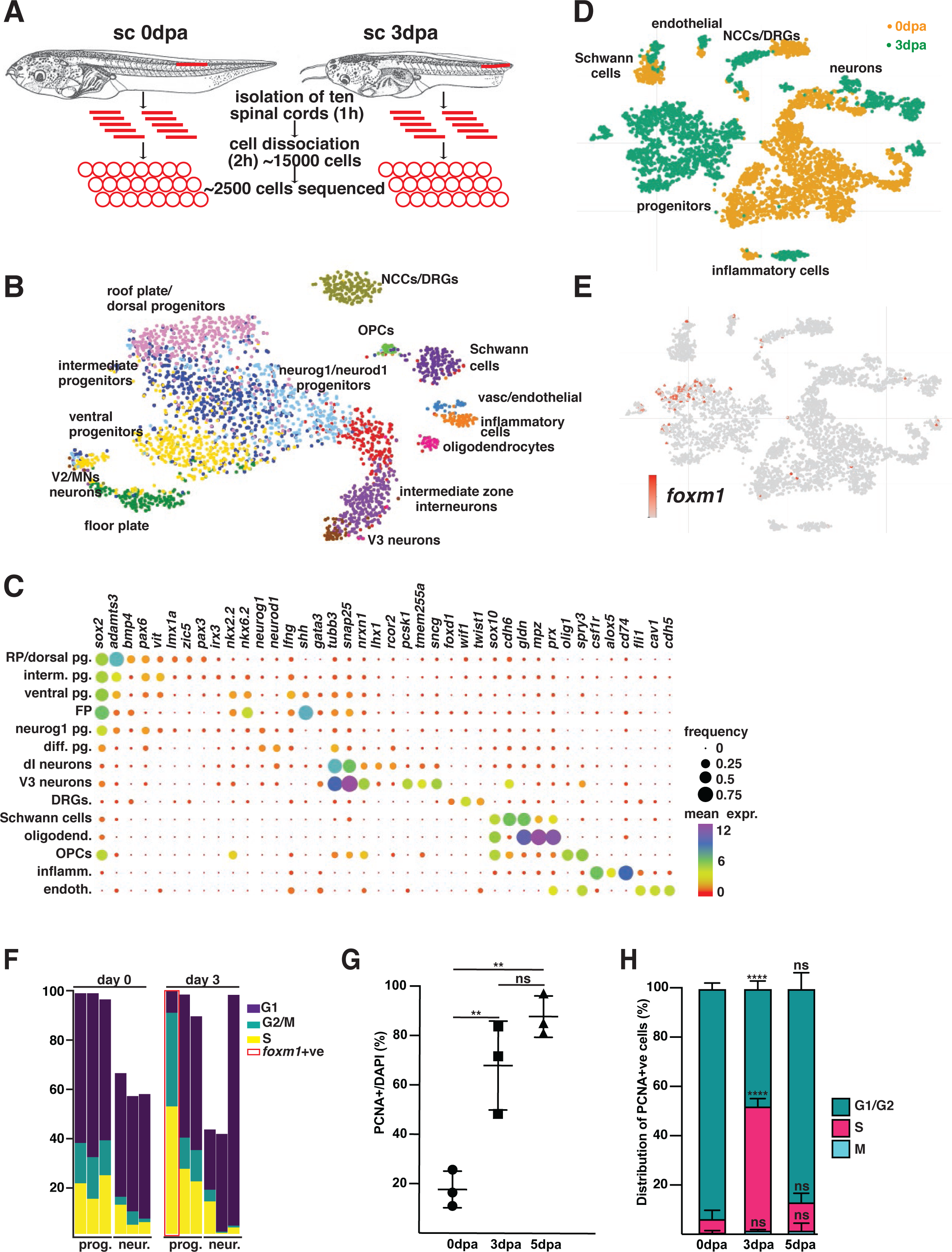
Single cell RNAseq (scRNAseq) identifies the characteristics of regenerating progenitors. **(A)** Schematic of the experimental design **(B)** t-SNE representation of the dataset from 0dpa with the different cell types identified **(C)** Bubble plot representing the percentage of cells (size of the dot) and level of expression (colour of the dot) for the genes used to identify the cell types in (B) **(D)** t-SNE representation of the whole dataset with cells from 0dpa in orange and from 3dpa in green **(E)** t-SNE plot showing the cells expressing *foxm1* **(F)** Bar plot showing the proportion of cells in G1, G2/M and S phase of clusters of progenitor cells (expressing *sox2* and *sox3*) and differentiated neurons (expressing *snap25b*). The total does not always amount to 100% as some clusters have cells from both 0 and 3dpa. The bar boxed in red represent the *foxm1* positive cluster. **(G)** The ratio of PCNA+ (left) per total number of cells (DAPI) in the spinal cord was determined at the indicated times after amputation (n=3, with at least 10 sections per data point). **(H)** The PCNA+ cells were then distributed in G1/G2 (diffuse signal), S (punctated signal) or M phase (condensed chromatin) at the indicated stage of regeneration. **p<0.01, ****p<0.0001, ns: non specific.

As the cellular organisation of the *Xenopus* spinal cord is not well described, we used the 10XGenomics platform to sequence 2503 cells from uninjured spinal cord. Fourteen clusters were identified comprising the different cell types expected in the spinal cord: roof and floor plate (*bmp4* and *shh* respectively), dorsal (*lmx1a/zic5)*, intermediate (*pax6, vit)* and ventral (*nkx2.2, nkx6.2)* progenitors, neurons (*tubb3, snap25)*, dorsal roots ganglia (*wif1* and *twist1*), oligodendrocytes (*mpz, prx)*, oligodendrocyte progenitor cells (OPCs; *sox2, olig1*), and Schwann cells (*sox10)*. Finally, we observed a small population of inflammatory (*csf1r, alox5*), vascular and endothelial (*fli1, cav1*) cells that may be spinal cord resident cells or a contamination from the dissection. However, no mesodermal or skin contamination was identified (Figures 3B and 3C).

We then analysed the changes in the transcriptome during regeneration (Figure 3D). A comparison of the clusters between 0dpa (yellow) and 3dpa (green) on a t-SNE representation shows that whilst some cell types cluster together (Schwann cells, DRGs), neural progenitors display a big shift in their transcriptome. Interestingly, *foxm1* is expressed in a specific cluster of neural progenitors only at 3dpa (Figure 3E). To characterise the *foxm1*+ cells, we identified DE genes between the *foxm1*+ cluster and the rest of the dataset (Figures S3B and S3C). The list of the 20 top DE genes ranked by false discovery rate (FDR) shows that the majority are upregulated and many are linked to the cell cycle (*ccna2, pcna, cdk2*). We then identified the GO terms that were significantly over-represented with PANTHER and used Revigo to generate a plot representation (Figure S3D). The majority of the GO terms identified are linked to the cell cycle. We therefore analysed changes in cell cycle dynamics between day0 and day3 (Figure 3F). Whilst clusters representing neurons are mainly in G1/G0 both at 0dpa and 3dpa, progenitor clusters have a higher proportion of cells in G2/M and S phases. Surprisingly, almost 50% of the cells in the *foxm1*+ cluster appears to be in S phase.

To confirm the changes in the cell cycle, we performed anti-PCNA staining on sections at 0, 3 and 5dpa (Figures 3G, 3H and S3E). The percentage of PCNA+ cells in the spinal cord increases sharply at 3dpa when compared to 0dpa (from 18 to 68%) and remains high until 5dpa (88%; Figure 3G). To estimate the proportion of cells in different phases of the cell cycle, we used the fact that PCNA expression is punctate in S phase and diffuse in G1/G2 phase [14,15] and the chromatin is condensed in M phase. We observed a transient increase of the cells in S phase from 6.6% at 0dpa to 45.7% at 3dpa with a return to baseline by 5dpa (12%; Figure 3H)). These data confirmed our scRNAseq experiments and indicate that proliferative cells in the spinal cord have a long S phase in early regeneration.

### Foxm1 regulates the fate of dividing progenitors during regeneration

Because of the changes in cell cycle and as Foxm1 promotes neuronal differentiation in early *Xenopus* development [16], we analysed the relative proportions of progenitors and neurons in the regenerating spinal cord when *foxm1* expression is impaired. Expression analysis by RT-qPCR shows an 33% increase (p=0.02) in *sox2* and a 18% (p=0.01) decrease in *ntub* expression in tadpoles with impaired *foxm1* expression (Figure 4A). Interestingly, reducing ROS levels with DPI also impairs expression of *ntub* and *ccnb3*, a well-characterised Foxm1 transcriptional target. However, DPI treatment had no effect on the expression of *ami*, a gene expressed in endothelial cells (Figure S4A).

**Figure 4.**
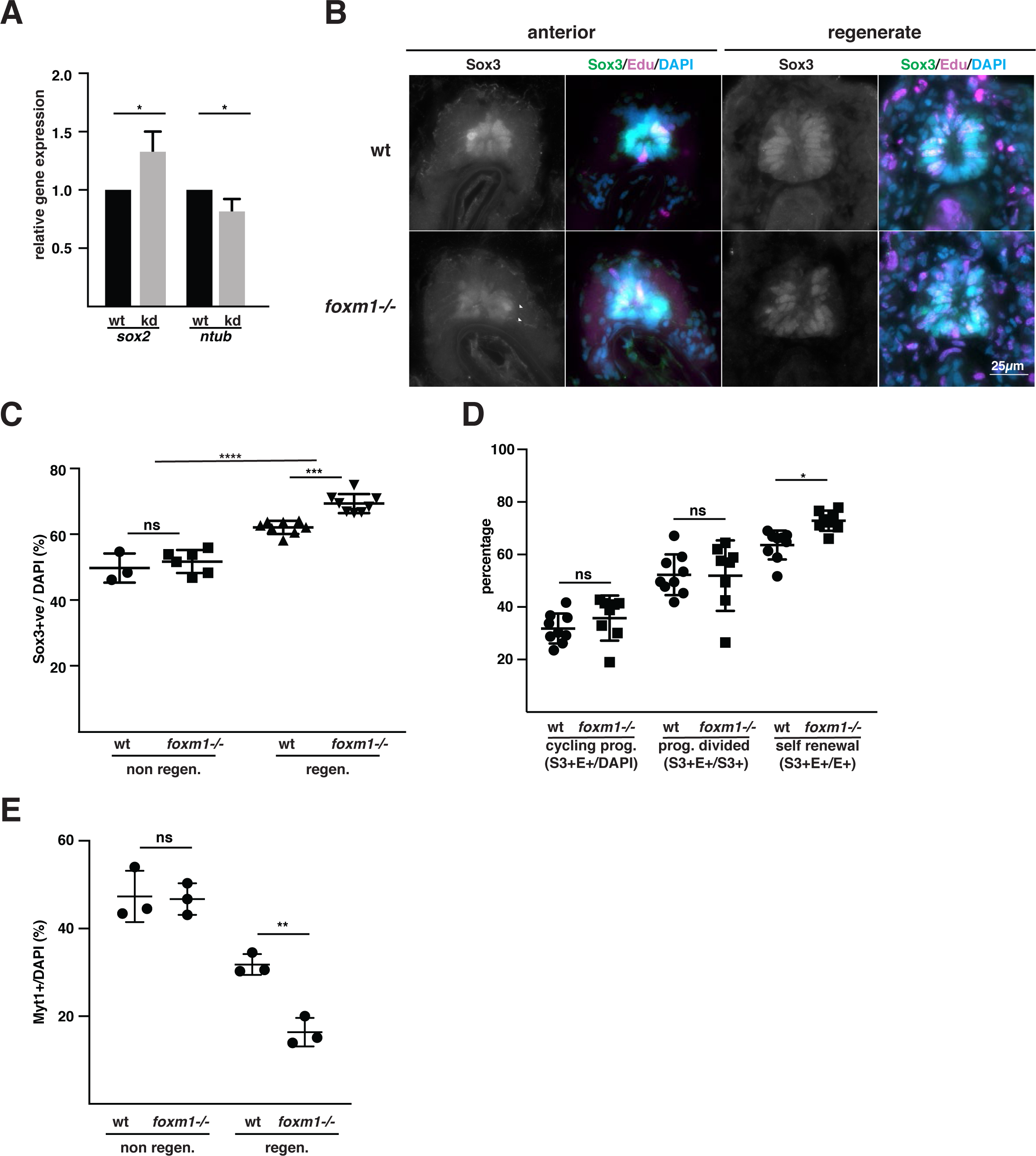
Foxm1 promotes the differentiation of neural progenitors in the regenerating spinal cord. **(A)** Tails from tadpoles *foxm1* knockdown (kd) and control (wt) NF50 tadpoles were amputated and left to regrow for 3 days. RNA was isolated from the regenerates and expression levels of *sox2* and *ntubulin* analysed by qPCR using *ef1*α as a reference. (*sox2*: n=4, *ntubulin*: n=6, with at least 20 tails per sample, mean +/-SD normalised to wt) **(B)** Representative images of sections from 5dpa regenerates sectioned and labelled with antibodies against Sox3 (green), EdU (red) and DAPI (blue). **(C)** Quantification of the images shown in (B). The ratio of Sox3+ (left) per total number of cells (DAPI) in the spinal cord was determined in the stump (non-regen.) and the regenerating spinal cord (regen.) in wt and *foxm1-/-* tadpoles (n=3 to 8, with at least 10 sections per data point). **(D)** Quantification of the proportion of cycling progenitors (Sox3+EdU+/DAPI, S3+/E+/DAPI), the proportion of progenitors having divided (Sox3+EdU+/Sox3+, S3+E+/S3+) and the proportion of progenitors self-renewal (Sox3+EdU+/EdU+, S3+E+/E+). **(E)** Tadpoles at stage NF50 were amputated and fixed 5 days later. The tails were then sectioned and stained using anti-Myt1 antibodies to label differentiated neurons (n=3 embryos with 10 sections each). For C, D and E, the graphs represent the mean +/- SD. ns=non-specific, *p<0.05, **p<0.01, ***p<0.001, ****p<0.0001.

We next analysed Sox3 expression by immunofluorescence using anti-Sox3 antibodies in the non-regenerating and regenerating spinal cord of wt and *foxm1*-/- NF50 tadpoles at 5dpa. As expected, in the non-regenerating spinal cord Sox3 is expressed in the cells lining the ventricle. We also observed Sox3+ extensions into the mantle zone of the spinal cord (Figure 4B, white arrowheads). In the wt regenerate, the spinal cord appears as an almost monolayer of cells around the central canal. Sox3 is expressed only in cells of the lateral spinal cord. These data reveal that the regenerating spinal cord conserved some cellular organisation. In contrast, in *foxm1*-/- tadpoles, we observed multiple cell layers and Sox3 expression is expressed more broadly (Figure 4B). To assess the distribution of cells in wt and *foxm1-/-* spinal cords, the angle of all nuclei (DAPI, Figures S4B and S4C) and Sox3+ nuclei (Figures S4D and S4E) relative to the centre of the central canal was calculated. Quantification of nuclei distribution either using cumulative number (Figure S4B) or normalised distribution (Figure S4C) show that whilst in the wt regenerating spinal cords, there are less cells laterally, this is not the case in *foxm1-/-* tadpoles. Furthermore, we observed a greater proportion of Sox3+ cells both dorsally and ventrally in the mutant compared to wt (Figures S4D and S4E).

The proportion of Sox3+ cells in the non-regenerating spinal cord is about 50% in wt and *foxm1*-/- tadpoles. In contrast, the proportion of progenitors increased in the regenerating spinal cord from 62.1+/-3.5% in wt to 69.4+/-2.9% in *foxm1-/-* (p<0.001, Figure 4C). Next, we labelled cycling cells with EdU at 3dpa and determined their fate at 5dpa using immunofluorescence. We first quantified the proportion of dividing progenitors (Edu+Sox3+) over the total number of cells (S3+E+/DAPI) or over the total number of progenitors (S3+E+/S3+, Figure 4D) tadpoles following amputation. Both ratios are similar in wt and *foxm1*-/- tadpoles, indicating that knocking out *foxm1* does not alter the rate of progenitor division. In contrast, the proportion of self-renewal (S3+E+/E+) increases significantly in the *foxm1-/-* compared to wt tadpoles (from 65% to 75%, p=0.006, Figure 4D). The stable rate of proliferation of progenitors combined with the increased rate in self renewal suggests that there is a shift in the fate of dividing progenitors from differentiation towards self-renewal.

To confirm these data, we analysed the expression of Myt1 as a neuronal marker in wt and *foxm1-/-* NF50 tadpoles at 5dpa (Figure 4E). Knocking out *foxm1* does not affect the percentage of Myt1+ cells in the non-regenerating spinal cord (∼43% of spinal cord cells). However, in the regenerate we observed a sharp reduction of Myt1+ cells in the mutant compared to wt (30% vs 14%, p<0.01). Similar data were obtained when we analysed F0 injected with gRNA targeting the *foxm1* locus compared to Cas9-injected controls (Figure S4F). Thus, in *foxm1* mutants, there is an increase of progenitors at the expense of differentiated neurons. Rather than promoting proliferation Foxm1 affects cell fate by promoting differentiation.

## Discussion

Using a combination of bulk and single-cell RNAseq experiments on isolated spinal cord during tail regeneration in *X.tropicalis*, we identified a new population of cells present exclusively in the regenerating spinal cord. It is characterised by the expression of *foxm1* and GO analysis reveals that genes involved in cell cycle and metabolism are over-represented. Impairing *foxm1* expression blocks spinal cord regeneration and alters the fate of the dividing progenitors with an increase in self-renewal division and a decrease in the production of neurons.

To characterise the *Xenopus* spinal cord at the molecular level, we undertook a single cell RNAseq approach. The number of cells and the depth of sequencing did not allow us to unambiguously determine the identity of progenitors and neurons subtypes but we identified the main cell types present in a vertebrate spinal cord, such as roof plate, dorsal and ventral progenitors, neurons and Schwann cells. Comparison of their transcriptomes at 0 and 3dpa reveals that differentiated cells did not respond much to injury. In contrast, the transcriptome progenitors displayed large changes as shown by their distinct locations on a t-SNE plot. Interestingly, at 3dpa, we isolate both the regenerate and the stump proximal to the amputation plan. However, no overlap of progenitors at 0dpa and 3dpa was observed, indicating that even cells positioned anteriorly from the amputation plan do respond to injury. Furthermore, *foxm1* expressing cells are a subset of the 3dpa progenitors, suggesting that this cluster represents the cells present in the regenerate.

Foxm1 has been shown to have a role in neuronal differentiation during primary neurogenesis in *Xenopus* [16] and in the mouse telencephalon [17]. However, in both cases, the role of Foxm1 seems to be dependent on its ability to control the overall length of the cell cycle. This is not the case in the regenerating spinal cord, where we do not observe a difference in proliferation in the regenerating spinal cord in *foxm1-/-* compared to controls. This suggests a cell-cycle independent role for Foxm1 during this process. Cell cycle regulators such as Cdc25b and Ccnd1 have been shown to play a role during neuronal development independently of their primary function [18–20]. Thus, it is possible that Foxm1 can promote neuronal differentiation in the regenerating spinal cord without affecting the overall length of the cell cycle.

Whilst we do not observe differences in the overall length of the cell cycle in mutant and wt tadpoles, a striking characteristic of the *foxm1+* cluster is the difference in the proportion of cells in each phase of the cell cycle compared to *foxm1-*progenitors. About 50% of the *foxm1*+ cells are in S-phase and 40% in G2/M, leaving only about 10% of cells in G1. Interestingly, similar changes in cell cycle have been observed in Axolotls, suggesting that it might be a general principle of spinal cord regeneration [8,21]. It has been suggested that long S phase might be necessary for progenitors undergoing self-renewal division to ensure genome integrity, especially in response to high level of ROS present in the nervous system [22–24]. The regenerating tail is an oxidative environment and we show here that *foxm1* expression requires ROS. Furthermore, Foxm1 has been shown to ensure chromosomal stability and genome integrity in U2OS and aged fibroblasts [25,26], raising the intriguing possibility that Foxm1 might ensure that the expansion of the progenitor pool does not lead to genetic instability during regeneration.

The expansion of the neural stem cell pool is required for spinal cord regeneration in axolotl, zebrafish and Xenopus [4,8,27]. Here we show that regeneration also requires the precise control of neuronal differentiation. In mammals, ependymal cells also re-enter the cell cycle upon spinal cord injury (SCI), are able to self-renew but differentiate mainly into astrocytes [28]. Understanding how Sox2/3+ cells are re-activated and able to differentiate into neurons may open new opportunity to enhance spinal cord regeneration in species that have limited regenerative capabilities.

## Material and Methods

### *Xenopus tropicalis* growth and tail amputation

*X.tropicalis* embryos were obtained and raised as previously described [29]. Tail amputation was performed at Nieuwkoop and Faber stages (NF)42-50 using a scalpel [30]. Tadpoles were anesthetized with 0.1% MS222 in 0.01X Marc’s Modified Ringer (MMR) solution (10mM NaCl, 0.2mM KCl, 0.1mM MgSO_4_, 0.2mM CaCl_2_, 0.5mM HEPES, pH 7.4) for the procedure followed by recovery in 0.01X MMR. All animal procedures complied with the UK Animal (Scientific Procedures) Act 1986 and were conducted with UK Home Office approval.

### RNA sequencing

Twenty spinal cords were isolated and immediately transferred into Trizol (Life Technologies) for each timepoint in triplicate. Following RNA extraction according to manufacturer’s instructions, total RNA concentration was quantified using Qubit HS RNA assay kit (Invitrogen) on the Qubit Fluorometer 2.0 (Invitrogen). Integrity was tested with the Agilent RNA 6000 Pico kit on the Agilent Bioanalyser. Samples with an RNA Integrity Number (RIN) ≥7 were considered of acceptable quality. RNAseq was performed with Illumina NextSeq500 using unpaired-end sequencing at the GeneCore facility (EMBL). Adapter sequences were trimmed using Trimmomatic v0.38. After quality control, the reads were converted to a FASTQ format and mapped on the v9.1 of the *X.tropicalis* transcriptome using bwa. The number of reads per transcript was determined using HTseq and the idxstats files were used to identify differentially expressed (DE) genes using the general linear model glmQLFit in DESeq2 with R. DE genes with a |log2(FC)|>1 and FDR<0.01 were considered significant. For the clustering, the k-mean was determined at 5 using the Elbow method and enriched gene ontology (GO) terms were identified using Fidea [31]. The full RNAseq dataset was then uploaded onto the Ingenuity Pathway Analysis (IPA, Qiagen) software. Upstream regulatory analysis was performed for DE genes with a |Log2FC|>1 and FDR<0.001 (992 genes for 0dpa/1dpa and 2720 genes for 0dpa/3dpa). This analysis predicts upstream molecules such as transcription factors that may be causing the observed change in gene expression [32]. RNA-seq data have been deposited in the ArrayExpress database at EMBL-EBI (www.ebi.ac.uk/arrayexpress) under accession number E-MTAB-8785.

For single cell RNAseq (scRNAseq), ten spinal cords at 0 and 3dpa were isolated and transferred in Modified Ringer’s (MR, 100mM NaCl, 1.8mM KCl, 2mM CaCl2, 1mM MgCl2, 5mM Hepes pH7) with 20µM Actinomycin D for 15 min to prevent *de novo* synthesis of mRNA [33]. The spinal cords were transferred in CMF-MR (100mM NaCl, 1.8mM KCl, 1mM EDTA and 5mM Hepes pH7), cut in small pieces using a fine scapel and incubated in 200µL of 0.5% Trypsin-EDTA without dye (GIBCO) for 30 min at 28°C with shaking at 600rpm. After a 5 min spin at 300rcf at 4°C, the spinal cords were incubated for 15 min at room temperature (RT) in 180µL of MR with 0.3U.µL^-1^ of DNaseI (D5025, Sigma). After trituration, 20µL of Collagenase IV at 50U.µL^-1^ was added and the samples were incubated 30 min at 28°C with shaking at 600rpm. The samples were passed through a flame elongated capillary, applied to a 5µm strainer and collected in MR with 0.0375% BSA. The samples were pelleted by centrifugation at 300rcf at 4°C for 5 min, washed with MR-BSA and resuspended in 35µL of MR-BSA. The cell number (4.10^5^ to 5.10^5^ cells.mL^-1^) and viability (above 90%) were estimated on a Countess Cell Counter (Invitrogen). The cells were then loaded onto the 10X Genomics platform for processing.

### Single cell RNAseq analysis

#### Building custom genome for mapping

The *X.tropicalis* genome v.9.1 was downloaded from Xenbase.org. A number of genes had duplicated entries where the same gene ID was assigned to different gene names. The majority of these genes had an overlapping transcript, we therefore used the GFFReads merging option to merge these transcripts.

#### Data pre-processing

The sequence files from the sequencer were processed using 10x Genomics custom pipeline Cell Ranger v2.2.0. The fastq files were aligned to the custom genome using the default parameters of Cell Ranger. The pipeline identified the barcodes associated with cells and counted UMIs mapped to each cell. Cell Ranger uses STAR aligner to align reads to the genome discarding all the counts mapping to multiple loci. The uniquely mapped UMI counts are reported in a gene by cell count matrix represented as a sparse matrix format. The Cell Ranger’s aggr command was used to aggregate the samples from 0 and 3dpa while keeping the default down-sampling parameter enabled.

#### Filtering

Low-quality cells were removed from the dataset to ensure that the technical noise did not affect the downstream analysis. Three commonly used parameters were used for cell quality evaluation: the number of UMIs per cell barcode (library size), the number of genes per cell barcode and the proportion of UMIs that are mapped to mitochondrial genes. Cells that have lower UMI counts than three Median Absolute Deviation (MAD) for the first two matrices and cells having higher proportion of reads mapped to mitochondrial genes with a cutoff of four MADs were filtered out.

After this initial filtering, 5271 cells (2503 0dpa and 2401 3dpa) out of 5411 cells remained for downstream analysis. Violin plots for these three metrices were then plotted to identify cells that have outlier distributions which can indicate doublets or multiplets of cells. However, no outliers were identified so no further filtering was done.

#### Classification of cell-cycle phase

Seurat’s CellCycleScoring method was used to calculate for each cell the score of S phase and G2M phase based on expression of S and G2/M phase markers. The cell cycle phase of each cell was identified based on the highest positive score of the phases. Cells that are expressing neither of the S phase and G2/M phase genes would have a negative value for both of these phases and were assigned as being in G1 [34].

#### Gene filtering and Normalization

Genes with average UMI counts below 0.01 were filtered out. After this filtering 11,215 genes were left for downstream analyses. To take into account the effect of variable library size for each cell, raw counts were normalized using the deconvolution-based method [35]. Counts from many cells are pooled together to circumvent the issue of higher number of zeros that are common in scRNA-seq data. This pool-based size-factors are then deconvoluted to find the size factor of each cell. These normalized data are then log-transformed with a pseudo-count of 1.

#### Visualization and Clustering

The first step for visualization and clustering is to identify the Highly Variable Genes (HVGs). To do this, we first decomposed the variance of each gene expression values into technical and biological components and identified the genes for which biological components were significantly greater than zero. These genes are called HVG. HVG genes were then used to reduce the dimensions of the dataset using PCA. The dimensions of dataset were further reduced to 2D using t -SNE, where 1 to 14 components of the PCA were given as input. The cells were grouped into their putative clusters using the dynamic tree cut method. Dynamic tree cut method identified the branch cutting point of a dendgoram dynamically and combined the advantage of both hierarchical and K-medoid clustering approach. This method identified 15 clusters in the population.

#### Identification of marker genes

To identify the marker gene for a cluster, we compared a given cluster with all other clusters. We selected the marker genes for the cluster using manual curation of DE genes with an FDR<0.01. For PANTHER analysis, all DE genes between the *foxm1*+ cluster were uploaded and tested against the pseudo-bulk of all the genes expressed in the scRNAseq experiment. scRNA-seq data have been deposited in the ArrayExpress database at EMBL-EBI (www.ebi.ac.uk/arrayexpress) under accession number E-MTAB-8839.

### CRISPR/Cas9 microinjection and inhibitor treatment

The *foxm1* CRISPR gRNA was designed using Crisprdirect (http://crispr.dbcls.jp) with the target sequence CCTGAGCAAACCCTTGTCCATGG. The gRNA cloning was carried out according to published protocols using the following primers fwd TAggAACTGTCAAGAAGGCGTTCC, rev AAACGGAACGCCTTCTTGACAGTT [36]. Eggs were injected with 300pg gRNA and 600ng Cas9 mRNA (from pT3TS-nCas9n, Addgene; Jao et al., 2013) or 600pg / 1.5ng of Cas9 Protein (M0386, NEB).

For chemical inhibitor treatments, tadpole tails were amputated at NF50 and left to recover for 36h in 0.01XMMR. Then, ROS signalling was inhibited with 4μM diphenyleneiodonium (DPI, Merck), FGF signalling with 20μM SU5402 (Calbiochem) and Shh signalling with 2.5μM of Cyclopamine (Merk). At 3dpa, the regenerates were collected, total RNA extracted and then processed for RT-qPCR.

### Genotyping

Genomic DNA was extracted by incubating the embryo or a section of the tail removed by amputation for 3h at 55°C in 10mM Tris pH8, 1mM EDTA, 80mM KCL, 0.3% NP40, 0.3% Triton X100 and 0.2mg.mL^-1^ of proteinase K. The samples were subsequently processed for PCR amplication using the following primers: fwd 5’-GTATGTTGCAGAGCAGGGCAT, rev 5’-GTATGTTGCAGAGCAGGGCAT. The PCR product was subsequently digested with NcoI for Restriction Fragment Length Polymorphism (RFLP) analysis.

### Isolation of RNA and qPCR

RNA was isolated from total embryos, tail regenerates and isolated spinal cord using Trizol (Life Technologies) according to manufacturer’s instructions. cDNA was generated using the Reverse Transcriptase AMV kit (Roche) for whole embryos and tails and Sensiscript Reverse Transcription Kit (Qiagen) for isolated spinal cords. qPCR analysis was performed on the StepOnePlus Real-Time PCR System using SYBR-Green reagents (Applied Biosystems). Expression was normalised to the expression levels of *ef1a* or *odc* and expression values were calculated using the ΔΔCt method.

The following primers were used (5’ to 3’ sequences): *foxm1*: fwd AAAGAGGAAGAGAGTGCGCC, rev TGGCATTTAGCTGCTCCTCC; *cyclinb3* fwd CTGCACTTCCACCATCCAATCCA, rev CAACTATATGCGGGACAGAGAG; *cdc25b* fwd GCCCAAACCCCTCGAGAAGA, rev GCCATCGAAGGTGCGTAGCCT; *ntubulin* fwd GGCAGTTACCATGGAGACAGT, rev GCCTGTGCCACCACCCAGAGA; *sox2* fwd CATGATGGAGACCGATCTCA, rev CTTACTCTGGTTGGAGCC; *ef1a* fwd GGATGGAACGGTGACAACATGCT, rev GCAGGGTAGTTCCGCTGCCAGA. The primers for *ami* and *odc* are described in [37]

### *In situ* hybridisation, EdU labelling and immunofluorescence

Whole mount in situ hybridisation on embryos and tails was performed as previously described [38,39]. The probe was generated from the TGas064p23 clone linearised with ClaI and transcribed using T7 polymerase.

For EdU labelling, NF50 tadpoles at 3dpa were injected with 3 times 4.2nL of 10mM EdU (Life Technologies) in DMSO for 3hr. After 2 days in normal media tails were collected and fixed in MEMFA. EdU was detected using Click-iT EdU Alexa Fluor 594 Imaging Kit (Life Technologies) following manufacturer’s instructions.

For immunostaining, the tails were fixed in MEMFA (0.1M MOPS, 5mM EDTA, 0.1M MgSO_4_, 4% formaldehyde), embedded in 25% fish gelatin / 20% sucrose, frozen on dry ice and sectioned at 12μm thickness on a Leica 3050S cryostat. The following antibodies were used: rabbit anti-Sox3 and anti-Myt1 were a gift from Nancy Papalopulu and used as described [40]; anti-PCNA was used at 1:500 (PC10, Sigma). Z stacks were acquired on a Cell Observer Z1 widefield microscope (Zeiss) using a 60X 1.4NA oil immersion objective. Images were then deconvolved using ZEN software (Adjustable deconvolution using fast iterative algorithm, Poisson (Richardson Lucy) likelihood, 40 iterations, 0.1% quality threshold). For the PCNA staining, the constrained iterative algorithm (Poisson (Richardson Lucy) likelihood, 40 iterations, 0.1% quality threshold) was used. Deconvolved maximum projections images were then used to quantify the different cell population using the cell counter module of Fiji.

### Statistics

When 2 or more conditions were compared, the normality of the distribution of the data was tested using a Shapiro-Wilk test. For two conditions, a two-tail unpaired t-test was used. If three or more conditions were compared, a one-way ANOVA followed by a Tukey Post-hoc test was used. For the rose plots and the cumulative distribution(Figure S4), a Kolmogorov-Smirnov test was used. All the statistical tests were done using Prism 8.

## Supplementary figure legends

**Figure S1 Metadata from the RNAseq experiment and regulation of *foxm1* expression (A)** Principle component analysis to assess the variation between all samples. The biological replicates of day 0 (0dpa, green), day 1 (1dpa, red) and day 3 post amputation (3dpa, blue) cluster together whilst showing wide variation in the two dimensions shown on the graph. **(B)** Hierarchical clustering of the nine datasets showing that the most important changes are observed between day 0 and day 3. **(C)** MA plot depicting the log fold change against the mean of normalised counts. DE genes (p_adj<0.05) are coloured in red. **(D)** Total number of differentially up- and down-regulated (p_adj<0.05) transcripts in 1dpa vs 0dpa and 3dpa vs 0dpa samples. (dpa = days post amputation). **(F)** Schematic of the experiment designed to establish what signals are upstream of *foxm1* expression. After amputation, the tails were left to heal for 36h before inhibitor treatments were started. The tails were collected at 72hpa and *foxm1* expression was determined by RT-qPCR. **(G)** Effects of treating tadpoles with 4*µ*M DPI (a NOX inhibitor) or DMSO as a control on *foxm1* expression. **(H, I)** Effects of inhibiting FGF signalling with 20*µ*M of SU5402 **(H)** and Hedgehog signalling with 2.5*µ*M Cyclopamin **(I)** on *foxm1* expression. The graphs represent the mean with SD of 4 independent experiments with about 15 tails per experiments. *p<0.05, ns: non specific

**Figure S2 Generation of a *foxm1* knockout line (A)** The CRISPR/Cas9 system was used to generate *foxm1* knockdown and knockout animals, gRNA was designed to target the *foxm1* gene. The target region contains the restriction site for NcoI and was used to test efficiency by RFLP. **(B)** Embryos were either uninjected (UI) or coinjected with gRNA and Cas9mRNA, 0.6ng Cas9 protein or 1.5ng Cas9 protein. Genomic DNA was extracted and a region amplified around the gRNA target site by PCR. Half of the PCR product was digested with NcoI. By comparing the ratio of the digested product with an intact restriction site (lower band) to the non-digested product containing a mutated restriction site (upper band) after the addition of NcoI (+) gives an indication of the efficiency of the induction of mutations. **(C)** Frogs injected with the CRISPR/Cas9 system and raised to adulthood. The F1 embryos were sequenced for mutations in *foxm1*. 2 frogs were identified to transmit frameshift mutations via the germline. **(D)** To test the efficiency of *foxm1* knockdown following CRISPR/Cas9 injection, injected embryos were raised to st24 and the RNA isolated. *foxm1* expression was analysed by qPCR, using *ef1*α as a reference (n=3 with 8 embryos per sample), data are expressed as the mean ± SD; *p=0.0251.

**Figure S3 Characterisation of the *foxm1* positive cells. (A)** Metadata of the single cell RNAseq experiment **(B)** Schematic representation of the cells used to identify DE genes and over-representation of GO terms. The blue cells correspond to the *foxm1* positive cluster. **(C)** Twenty most significantly DE genes comparing the blue versus the red cells ranked by FDR. **(D)** GO-Slim Biological Process terms over-represented were identified by uploading the DE genes into PANTHER. The GO terms significantly upregulated were then inputted into Revigo (http://revigo.irb.hr/) to generate a plot representation. **(E)** Representative section labelled with an anti-PCNA antibody at the indicated day after amputation (dpa). The white arrowheads point to cells in S phase in the spinal cord and the purple arrowhead at cells in S phase in the surrounding tissue.

**Figure S4 Effect of impairing *foxm1* expression on the organisation of the regenerating spinal cord. (A)** Tails of tadpoles at stage NF50 were amputated and 36h later treated with 4*µ*M DPI or DMSO as a control until 72hpa. Total RNA was isolated from regenerating tails, reversed transcribed into cDNA and analysed for the indicated transcript expression by qPCR, using *ef1*α as a reference gene (n=3, the graph represents the mean with SD normalised to control). **(B)** Rose plot histograms showing the spread of cells in the wt (blue) or *foxm1*-/- (red) spinal cord, presented as angles from the centre of the central canal. **(C)** Percentage frequency distribution of the angles of DAPI+ nuclei in wt (blue) or *foxm1*-/- spinal cords (red). p<0.0001 as analysed by Kolmogorov-Smirnov tests. **(D)** Rose plot histograms showing the spread of Sox3+ cells in the wt (blue) or *foxm1*-/- (red) spinal cord, presented as angles from the centre of the central canal. **(E)** Percentage frequency distribution of the angles of DAPI+ nuclei in wt (blue) or *foxm1*-/- spinal cords (red). p<0.0001 as analysed by Kolmogorov-Smirnov tests. (A-D) angles are distributed into 10 bins from 0 – 180 degrees using a Matlab script. Dorsal = 0 degrees, lateral = 90 degrees, ventral = 180 degrees. Ten sections from n = 2 animals were analysed per genotype. Total cell counts were as follows: wt (DAPI+, 878 nuclei; Sox3+, 397), foxm1-/- (DAPI+, 1233; Sox3+, 658). **(F)** The tails of control and *foxm1*KD were amputated and 5 days post amputation tails were sectioned and labelled with the Sox3 or Myt1 antibody followed by DAPI staining. The ratio of Sox3 (left panel) and Myt1 (right panel) per number of DAPI stained nuclei in the spinal cord was quantified and compared between control and foxm1KD tadpoles. (Sox3 wt n=6 with 41 sections, CRISPR/Cas9 n=5 with 50 sections; Myt1 wt = 8 with 67 sections, foxm1KD n=7 with 53 sections). The graph shows the mean +/- SD and the significance was tested by ANOVA with a Tukey post-hoc test (***p=0.0006; ****p<0.0001).

## Acknowledgments

DP and LP are supported by a BBSRC-DTP PhD studentships. This work was supported by a Wellcome Trust Seed Award (ref 205894/Z/17/Z), the Manchester Regenerative Medicine Network (MaRMN), The University of Manchester Strategic Fund (ISSF) to K.D and MRC single-cell centre award [MR/M008908/1] to S.M.B. We thank the EMBL core facility for performing the bulk RNAseq experiments, the Genomics Core Facility (University of Manchester) for the scRNAseq and the National Xenopus Resource (NXR, Marine Biology Laboratories, Woods Hole, USA) for their bioinformatics workshop. We thank Nancy Papalopulu (University of Manchester) for the anti-Sox3 and anti-Myt1 antibodies and Enrique Amaya, Raman Das and Shane Herbert for comments on the manuscript.

## Author contributions

Conceptualisation, D.P. and K.D.; Methodology, D.P., L.P., S.M.B. and K.D.; Software, L.P., S.M.B. and K.D.; Investigation, D.P., L.P., R.T. and K.D.; Data curation, S.M.B; Writing – original draft, K.D.; Review & editing, D.P., L.P., R.T., S.M.B. and K.D.; Supervision, K.D.; Funding acquisition, K.D.

## References

1. McDonald, J.W., and Sadowsky, C. (2002). Spinal-cord injury. Lancet 359, 417–425.

2. Slack, J.M.W., Lin, G., and Chen, Y. (2008). The Xenopus tadpole: a new model for regeneration research. Cell. Mol. Life Sci. 65, 54–63.

3. Love, N.R., Chen, Y., Bonev, B., Gilchrist, M.J., Fairclough, L., Lea, R., Mohun, T.J., Paredes, R., Zeef, L.A.H., and Amaya, E. (2011). Genome-wide analysis of gene expression during Xenopus tropicalis tadpole tail regeneration. BMC Dev Biol 11, 70.

4. Muñoz, R., Edwards-Faret, G., Moreno, M., Zuñiga, N., Cline, H., and Larraín, J. (2015). Regeneration of Xenopus laevis spinal cord requires Sox2/3 expressing cells. Dev. Biol. 408, 229–243.

5. Gaete, M., Muñoz, R., Sánchez, N., Tampe, R., Moreno, M., Contreras, E.G., Lee-Liu, D., and Larraín, J. (2012). Spinal cord regeneration in Xenopus tadpoles proceeds through activation of Sox2-positive cells. Neural Dev 7, 13.

6. McHedlishvili, L., Epperlein, H.H., Telzerow, A., and Tanaka, E.M. (2007). A clonal analysis of neural progenitors during axolotl spinal cord regeneration reveals evidence for both spatially restricted and multipotent progenitors. Development 134, 2083–2093.

7. Rost, F., Rodrigo Albors, A., Mazurov, V., Brusch, L., Deutsch, A., Tanaka, E.M., and Chara, O. (2016). Accelerated cell divisions drive the outgrowth of the regenerating spinal cord in axolotls. Elife 5.

8. Rodrigo Albors, A., Tazaki, A., Rost, F., Nowoshilow, S., Chara, O., and Tanaka, E.M. (2015). Planar cell polarity-mediated induction of neural stem cell expansion during axolotl spinal cord regeneration. Elife 4, e10230.

9. Chang, J., Baker, J., and Wills, A. (2017). Transcriptional dynamics of tail regeneration in Xenopus tropicalis. Genesis 55, e23015.

10. Love, N.R., Chen, Y., Ishibashi, S., Kritsiligkou, P., Lea, R., Koh, Y., Gallop, J.L., Dorey, K., and Amaya, E. (2013). Amputation-induced reactive oxygen species are required for successful Xenopus tadpole tail regeneration. Nat Cell Biol 15, 222–228.

11. Lin, G., and Slack, J.M. (2008). Requirement for Wnt and FGF signaling in Xenopus tadpole tail regeneration. Dev. Biol. 316, 323–335.

12. Beck, C.W., Christen, B., and Slack, J.M. (2003). Molecular pathways needed for regeneration of spinal cord and muscle in a vertebrate. Dev. Cell 5, 429–439.

13. Schüller, U., Zhao, Q., Godinho, S.A., Heine, V.M., Medema, R.H., Pellman, D., and Rowitch, D.H. (2007). Forkhead transcription factor FoxM1 regulates mitotic entry and prevents spindle defects in cerebellar granule neuron precursors. Mol Cell Biol 27, 8259–8270.

14. Rottach, A., Kremmer, E., Nowak, D., Boisguerin, P., Volkmer, R., Cardoso, M.C., Leonhardt, H., and Rothbauer, U. (2008). Generation and Characterization of a Rat Monoclonal Antibody Specific for PCNA. Hybridoma 27, 91–98.

15. Celis, J.E., Madsen, P., Nielsen, S., and Celis, A. (1986). Nuclear patterns of cyclin (PCNA) antigen distribution subdivide S-phase in cultured cells — Some applications of PCNA antibodies. Leuk. Res. 10, 237–249.

16. Ueno, H., Nakajo, N., Watanabe, M., Isoda, M., and Sagata, N. (2008). FoxM1-driven cell division is required for neuronal differentiation in early Xenopus embryos. Development 135, 2023–2030.

17. Wu, X., Gu, X., Han, X., Du, A., Jiang, Y., Zhang, X., Wang, Y., Cao, G., and Zhao, C. (2014). A novel function for Foxm1 in interkinetic nuclear migration in the developing telencephalon and anxiety-related behavior. J. Neurosci. 34, 1510–1522.

18. Hydbring, P., Malumbres, M., and Sicinski, P. (2016). Non-canonical functions of cell cycle cyclins and cyclin-dependent kinases. Nat Rev Mol Cell Biol 17, 280–292.

19. Bonnet, F., Molina, A., Roussat, M., Azais, M., Vialar, S., Gautrais, J., Pituello, F., and Agius, E. (2018). Neurogenic decisions require a cell cycle independent function of the CDC25B phosphatase. Elife 7, e32937.

20. Lukaszewicza, A.I., and Anderson, D.J. (2011). Cyclin D1 promotes neurogenesis in the developing spinal cord in a cell cycle-independent manner. Proc. Natl. Acad. Sci. U. S. A. 108, 11632–11637.

21. Costa, E.C., Albors, A.R., Tanaka, E.M., and Chara, O. (2020). Modeling the spatiotemporal control of cell cycle acceleration during axolotl spinal cord regeneration. bioRxiv, 2020.02.10.941443.

22. Arai, Y., Pulvers, J.N., Haffner, C., Schilling, B., Nüsslein, I., Calegari, F., and Huttner, W.B. (2011). Neural stem and progenitor cells shorten S-phase on commitment to neuron production. Nat. Commun. 2, 154.

23. Turrero García, M., Chang, Y., Arai, Y., and Huttner, W.B. (2016). S-phase duration is the main target of cell cycle regulation in neural progenitors of developing ferret neocortex. J. Comp. Neurol. 524, 456–470.

24. Narciso, L., Parlanti, E., Racaniello, M., Simonelli, V., Cardinale, A., Merlo, D., and Dogliotti, E. (2016). The Response to Oxidative DNA Damage in Neurons: Mechanisms and Disease. Neural Plast. 2016, 1–14.

25. Laoukili, J., Kooistra, M.R.H., Brás, A., Kauw, J., Kerkhoven, R.M., Morrison, A., Clevers, H., and Medema, R.H. (2005). FoxM1 is required for execution of the mitotic programme and chromosome stability. Nat Cell Biol 7, 126–136.

26. Macedo, J.C., Vaz, S., Bakker, B., Ribeiro, R., Bakker, P.L., Escandell, J.M., Ferreira, M.G., Medema, R., Foijer, F., and Logarinho, E. (2018). FoxM1 repression during human aging leads to mitotic decline and aneuploidy-driven full senescence. Nat. Commun. 9.

27. Ogai, K., Nakatani, K., Hisano, S., Sugitani, K., Koriyama, Y., and Kato, S. (2014). Function of Sox2 in ependymal cells of lesioned spinal cords in adult zebrafish. Neurosci Res 88, 84–87.

28. Meletis, K., Barnabé-Heider, F., Carlén, M., Evergren, E., Tomilin, N., Shupliakov, O., and Frisén, J. (2008). Spinal cord injury reveals multilineage differentiation of ependymal cells. PLoS Biol. 6, 1494–1507.

29. Collu, G.M., Hidalgo-Sastre, A., Acar, A., Bayston, L., Gildea, C., Leverentz, M.K., Mills, C.G., Owens, T.W., Meurette, O., Dorey, K., et al. (2012). Dishevelled limits Notch signalling through inhibition of CSL. Development 139, 4405–4415.

30. Nieuwkoop, P.D., and Faber, J. (1994). Normal table of Xenopus laevis (Daudin) : a systematical and chronological survey of the development from the fertilized egg till the end of metamorphosis.

31. D’Andrea, D., Grassi, L., Mazzapioda, M., and Tramontano, A. (2013). FIDEA: a server for the functional interpretation of differential expression analysis. Nucleic Acids Res. 41, W84–W88.

32. Krämer, A., Green, J., Pollard, J., and Tugendreich, S. (2014). Causal analysis approaches in ingenuity pathway analysis. Bioinformatics 30, 523–530.

33. Wu, Y.E., Pan, L., Zuo, Y., Li, X., and Hong, W. (2017). Detecting Activated Cell Populations Using Single-Cell RNA-Seq. Neuron 96, 313–329.

34. Aztekin, C., Hiscock, T.W., Marioni, J.C., Gurdon, J.B., Simons, B.D., and Jullien, J. (2019). Identification of a regeneration-organizing cell in the Xenopus tail. Science (80-.). 364, 653–658.

35. Lun, A.T.L., Bach, K., and Marioni, J.C. (2016). Pooling across cells to normalize single-cell RNA sequencing data with many zero counts. Genome Biol. 17, 75.

36. Jao, L.-E., Wente, S.R., and Chen, W. (2013). Efficient multiplex biallelic zebrafish genome editing using a CRISPR nuclease system. Proc Natl Acad Sci U S A 110, 13904–13909.

37. Nagamori, Y., Roberts, S., Maciej, M., and Dorey, K. (2014). Activin ligands are required for the re-activation of Smad2 signalling after neurulation and vascular development in Xenopus tropicalis. Int. J. Dev. Biol. 58, 783–791.

38. Harland, R.M. (1991). In situ hybridization: an improved whole-mount method for Xenopus embryos. Methods Cell Biol. 36, 685–695.

39. Lea, R., Papalopulu, N., Amaya, E., and Dorey, K. (2009). Temporal and spatial expression of FGF ligands and receptors during Xenopus development. Dev Dyn 238, 1467–1479.

40. Thuret, R., Auger, H., and Papalopulu, N. (2015). Analysis of neural progenitors from embryogenesis to juvenile adult in Xenopus laevis reveals biphasic neurogenesis and continuous lengthening of the cell cycle. Biol. Open 4, 1772–1781.

