## Supplementary figures and images for "Foxm1 regulates neuronal progenitor fate during spinal cord regeneration"

### FIgure S1

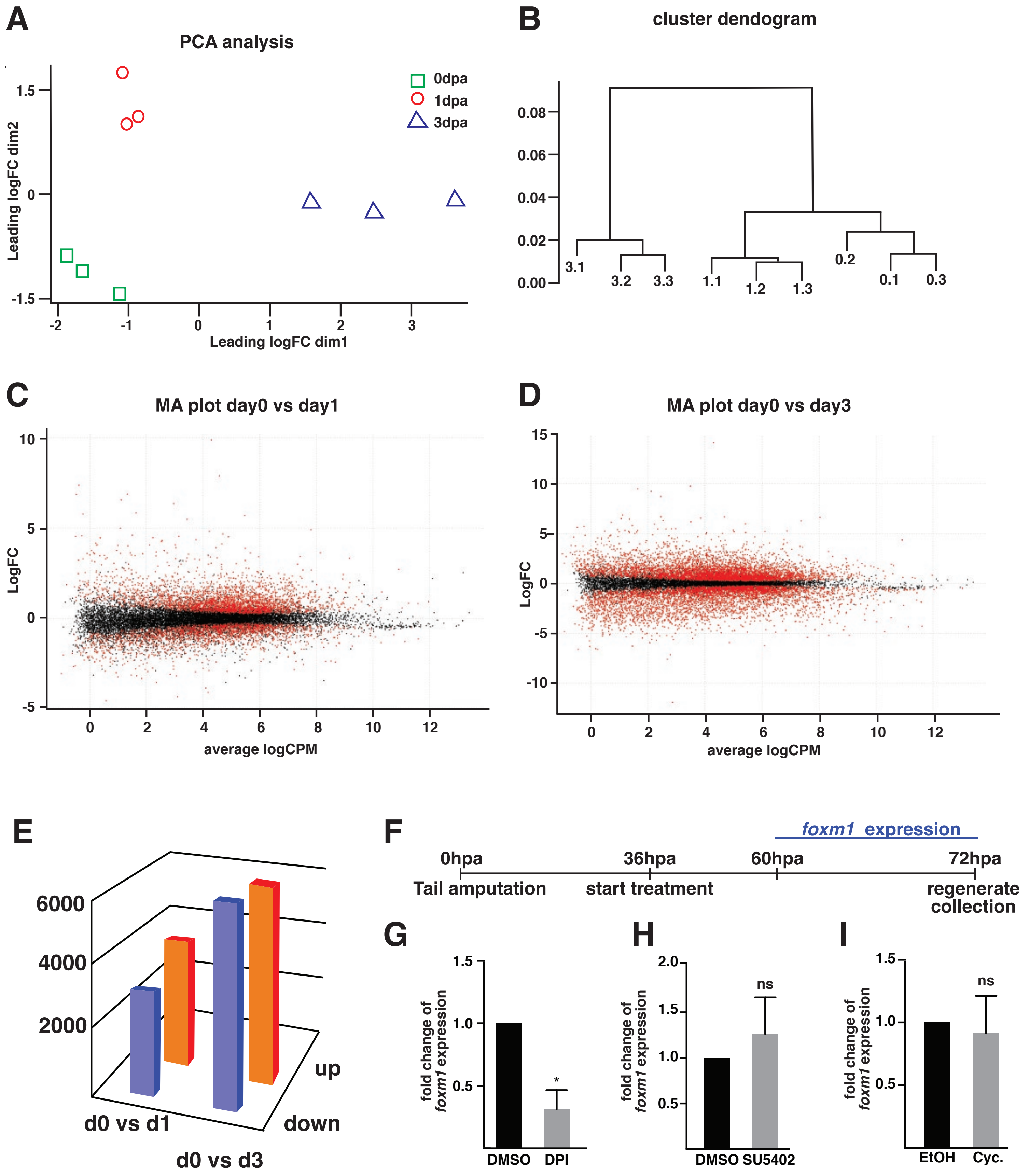


Figure S1 Pelzer *et al.*

### Figure S2

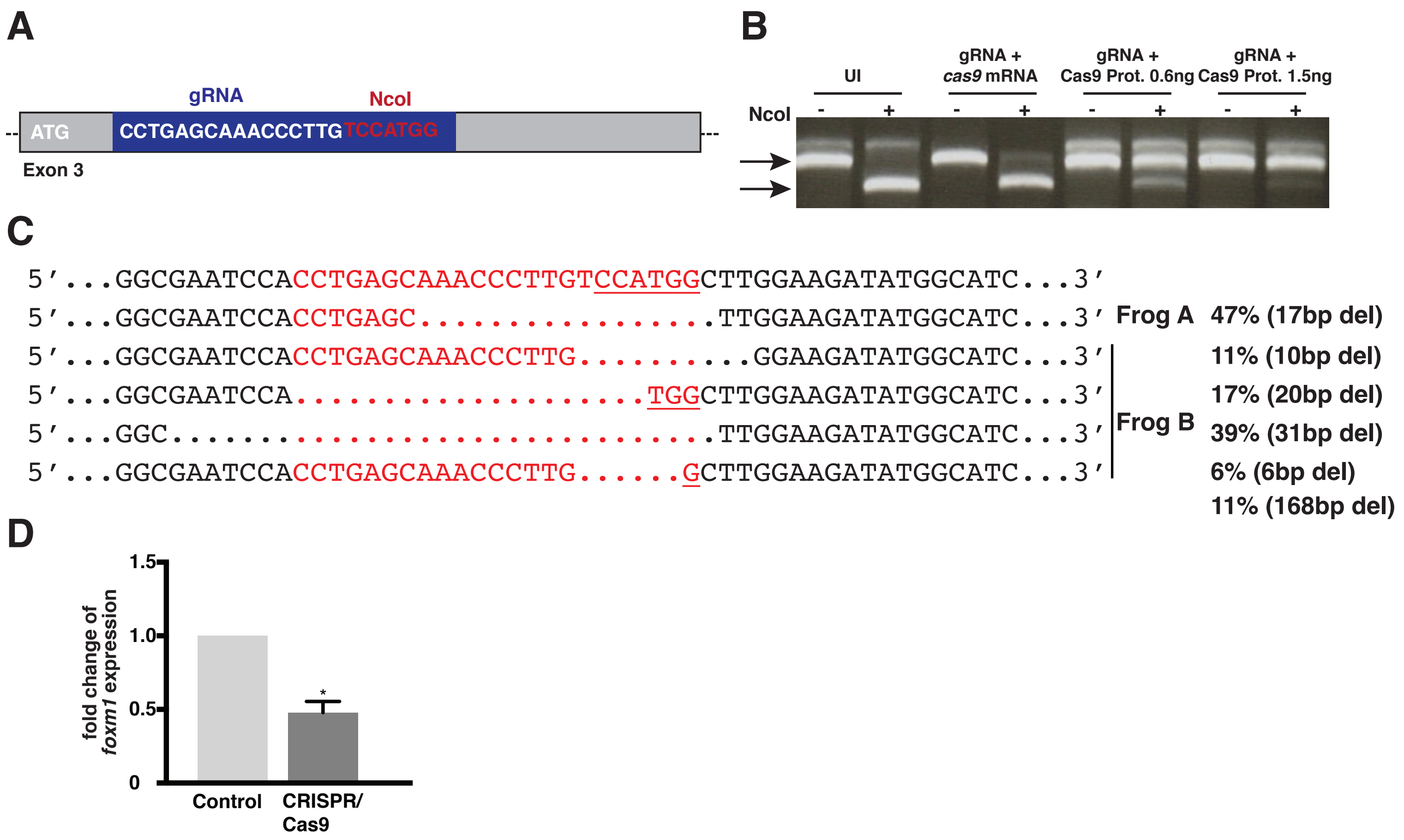


Figure S2 Pelzer *et al.*

### Figure S4

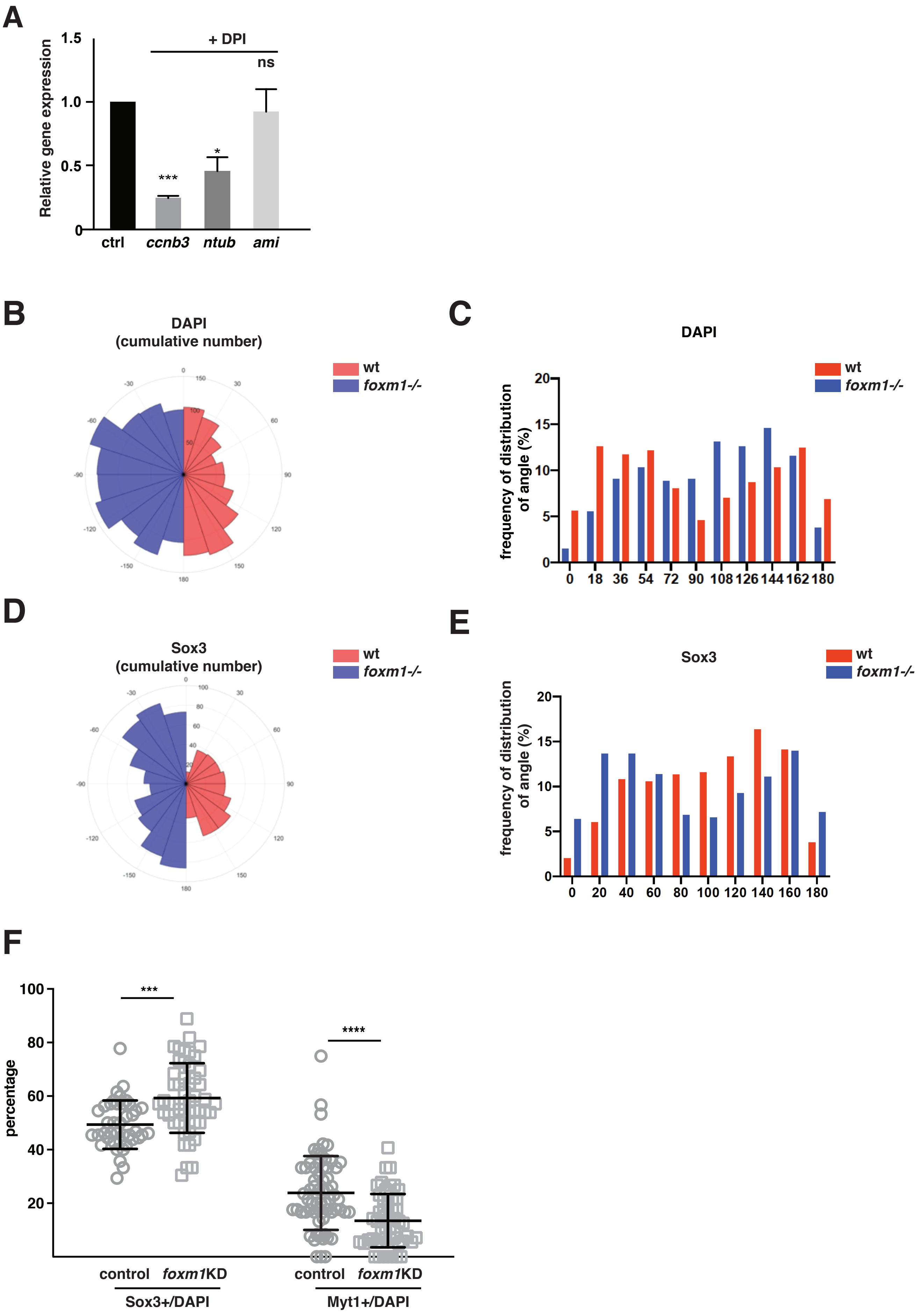


**Figure S4 Pelzer *et al.***
