## Supplementary material for "Foxm1 regulates neuronal progenitor fate during spinal cord regeneration": Figure S3

**A**

| Sample ID | 0dpa | 3dpa |
|---|---|---|
| Stage | 50 | 50 |
| Days post amputation | 0 | 3 |
| Sequencing Batch | 1 | 1 |
| Number of cells | 2503 | 2401 |
| Mean reads per cell | 75,815 | 87,857 |
| Median genes per cell | 590 | 1548 |
| Median UMI counts per cell | 2,679 | 7,109 |
| Total genes detected | 16,839 | 18.87 |
| Fraction reads in cells (%) | 88 | 88.5 |

**B**

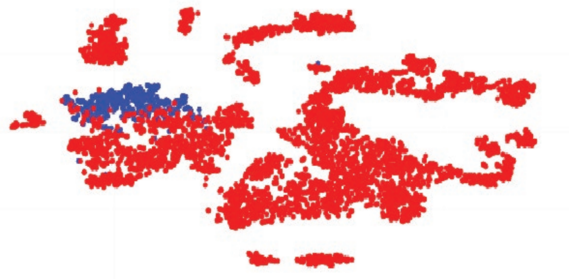

**C**

| Gene name | Log2(FC) | FDR |
|---|---|---|
| rsph9 | 4.06 | 3.35E-106 |
| ccna2 | 4.89 | 2.33E-105 |
| pcna | 3.99 | 1.36E-102 |
| rrm2.2 | 4.34 | 6.98E-100 |
| tubb4b | 3.85 | 3.18E-99 |
| cks2 | 4.38 | 2.03E-95 |
| birc5.2 | 4.40 | 2.71E-92 |
| kiaa0101 | 3.86 | 8.12E-90 |
| hmgb2 | 3.55 | 1.20E-88 |
| odf3 | 4.43 | 2.05E-85 |
| mcm6.2 | 4.12 | 9.70E-85 |
| stmn2 | -4.75 | 7.00E-84 |
| LOC100498440 | 3.96 | 2.77E-82 |
| incenp | 4.53 | 2.27E-80 |
| nme5 | 3.87 | 5.62E-80 |
| mcm5 | 3.98 | 5.21E-76 |
| tuba3c | 3.64 | 9.35E-75 |
| cdk2 | 3.74 | 1.17E-74 |
| kif20b | 4.22 | 3.11E-74 |
| mcm7 | 3.81 | 9.72E-73 |
| cdca7 | 3.77 | 1.13E-72 |

**D**

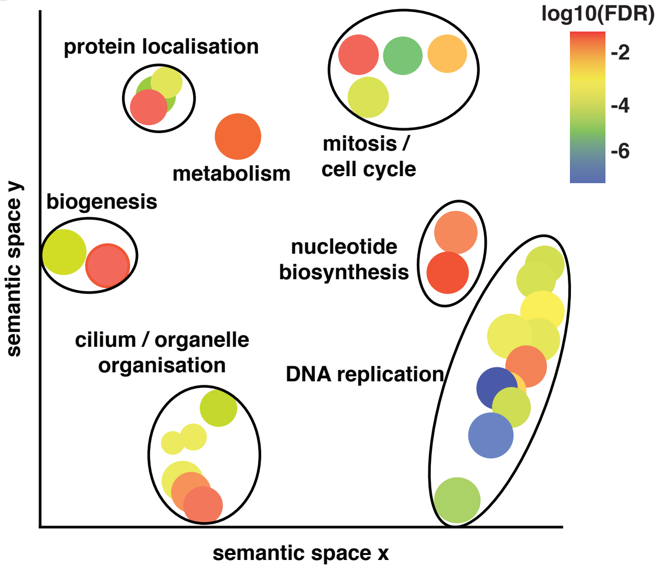


**E**

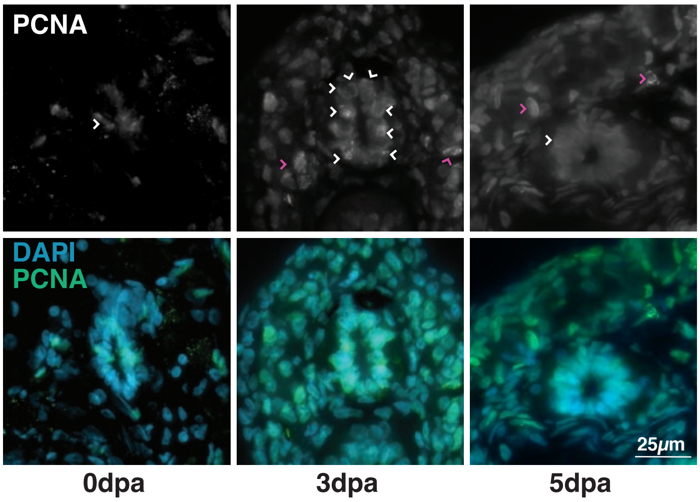


**Figure S3 Pelzer *et al.***
